# A transcriptomic atlas of mammalian olfactory mucosae reveals an evolutionary influence on food odor detection in humans

**DOI:** 10.1101/552232

**Authors:** Luis R. Saraiva, Fernando Riveros-McKay, Massimo Mezzavilla, Eman H. Abou-Moussa, Charles J. Arayata, Casey Trimmer, Ximena Ibarra-Soria, Mona Khan, Laura Van Gerven, Mark Jorissen, Matthew Gibbs, Ciaran O’Flynn, Scott McGrane, Peter Mombaerts, John C. Marioni, Joel D. Mainland, Darren W. Logan

**Affiliations:** Sidra Medicine, PO Box 26999, Doha, Qatar; Wellcome Sanger Institute, Wellcome Genome Campus, Hinxton-Cambridge, CB10 1SD, United Kingdom; European Bioinformatics Institute (EMBL-EBI), European Molecular Biology Laboratory, Wellcome Genome Campus, Hinxton-Cambridge, CB10 1SD, United Kingdom; Monell Chemical Senses Center, Philadelphia, PA 19104, USA; Max Planck Research Unit for Neurogenetics, Max von-Laue-Strasse 4, 60438 Frankfurt, Germany; Department of ENT-HNS, UZ Leuven, Herestraat 49, 3000 Leuven, Belgium; Waltham Centre for Pet Nutrition, Leicestershire, LE14 4RT, UK; Department of Neuroscience, University of Pennsylvania, Philadelphia, PA, 19104, USA

## Abstract

The mammalian olfactory system displays species-specific adaptations to different ecological niches. To investigate the evolutionary dynamics of olfactory sensory neuron (OSN) sub-types across 95 million years of mammalian evolution, we applied RNA-sequencing of whole olfactory mucosa samples from mouse, rat, dog, marmoset, macaque and human. We find that OSN subtypes representative of all known mouse chemosensory receptor gene families are present in all analyzed species. Further, we show that OSN subtypes expressing canonical olfactory receptors (ORs) are distributed across a large dynamic range and that homologous subtypes can be either highly abundant across all species or species/order-specific. Interestingly, highly abundant mouse and human OSN subtypes detect odorants with similar sensory profiles, and sense ecologically relevant odorants, such as mouse semiochemicals or human key food odorants. Taken together, our results allow for a better understanding of the evolution of mammalian olfaction in mammals and provide insights into the possible functions of highly abundant OSN subtypes in mouse and human.

## MAIN TEXT

Odor detection in mammals is initiated by the activation of olfactory receptors (ORs) expressed in olfactory sensory neurons (OSNs), which populate the whole olfactory mucosa (WOM) (Buck and Axel, 1991). Most mature OSNs predominantly express one allele of a single OR gene (Chess et al., 1994; Hanchate et al., 2015; Malnic et al., 1999; Saraiva et al., 2015b). Smaller subsets of mature OSNs express other families of chemosensors, such as trace-amine associated receptors (TAARs), guanylate cyclases (GCs), or members of the membrane-spanning 4-pass A (MS4As) gene family (Fulle et al., 1995; Greer et al., 2016; Hussain et al., 2009; Liberles and Buck, 2006; Omura and Mombaerts, 2015; Saraiva et al., 2015b). These receptors define the molecular identity and odorant response profile of OSNs, and OSNs apply a combinatorial strategy to discriminate a number of odorants vastly greater than the number of receptors present in the genome (Malnic et al., 1999; Nara et al., 2011; Zhang et al., 2013). From a phylogenetic perspective, OR genes are divided in two classes: class I, which preferentially bind hydrophilic odorants, and class II, which tend to recognize hydrophobic odorants (Saito et al., 2009; Zhang and Firestein, 2002). The complex evolutionary dynamics of OR genes have resulted in strikingly different species-specific repertoires, which are presumably shaped by the chemosensory information that is required for survival in each species’ niche (Bear et al., 2016; Khan et al., 2015; Nei et al., 2008; Niimura, 2012; Niimura et al., 2014; Niimura and Nei, 2005). While cataloging the presence of orthologous OR genes among species has provided some insight into the drivers of selection (Niimura et al., 2014), knowing the relative abundance of each OSN subtype both within and among species may provide a better understanding of these evolutionary dynamics.

Using an RNA-sequencing (RNA-seq) based approach, we have profiled previously the complete mouse and zebrafish OSN repertoires and found that they are stratified into hundreds to thousands of functionally distinct subtypes, represented across a large dynamic range of abundance in both species (Ibarra-Soria et al., 2014; Saraiva et al., 2015a; Saraiva et al., 2015b). While this OSN distribution is stereotyped among genetically identical mice, it varies greatly among different strains (Ibarra-Soria et al., 2017). These distributions are largely genetically controlled in *cis* and have thus likely diverged under evolutionary pressures (Ibarra-Soria et al., 2017). The abundance of OSNs expressing a given OR correlates with the total volume of corresponding glomeruli in the olfactory bulb (Bressel et al., 2016). Increasing the number of OR-expressing OSNs in mouse lowers detection thresholds (D’Hulst et al., 2016), raising the possibility that more abundant expression increases sensitivity to the receptor’s ligands, providing a mechanism for adaptation to enhance the detection of important ecological olfactory cues. Therefore, olfactory transcriptome analysis is a critical first step towards identifying the most functionally significant among the hundreds of OSN subtypes without identified odorants. Here, we investigated the transcriptional dynamics and putative functions of the olfactory systems of six mammalian species, spanning ∼95 million years of evolution.

We performed RNA-seq on the WOM of male dog (*Canis familiaris*), mouse (*Mus musculus*), rat (*Rattus norvegicus*), marmoset (*Callithrix jacchus*), macaque (*Macaca mulatta*), and human (*Homo sapiens*) (Fig. 1A). We processed three biological replicates for each species, except for macaque, where we profiled four (File S1 for quality metrics, File S2 for all gene expression data). On average, 83.85 ± 1.66 % of the total reads mapped uniquely to each corresponding genome. The intraspecific variability level among WOM replicates is extremely low (spearman’s rho, r_s_=0.95-0.98, P<0.0001), consistent with previous studies in laboratory animals (Ibarra-Soria et al., 2014; Ibarra-Soria et al., 2017; Saraiva et al., 2015a).

**Figure 1.**
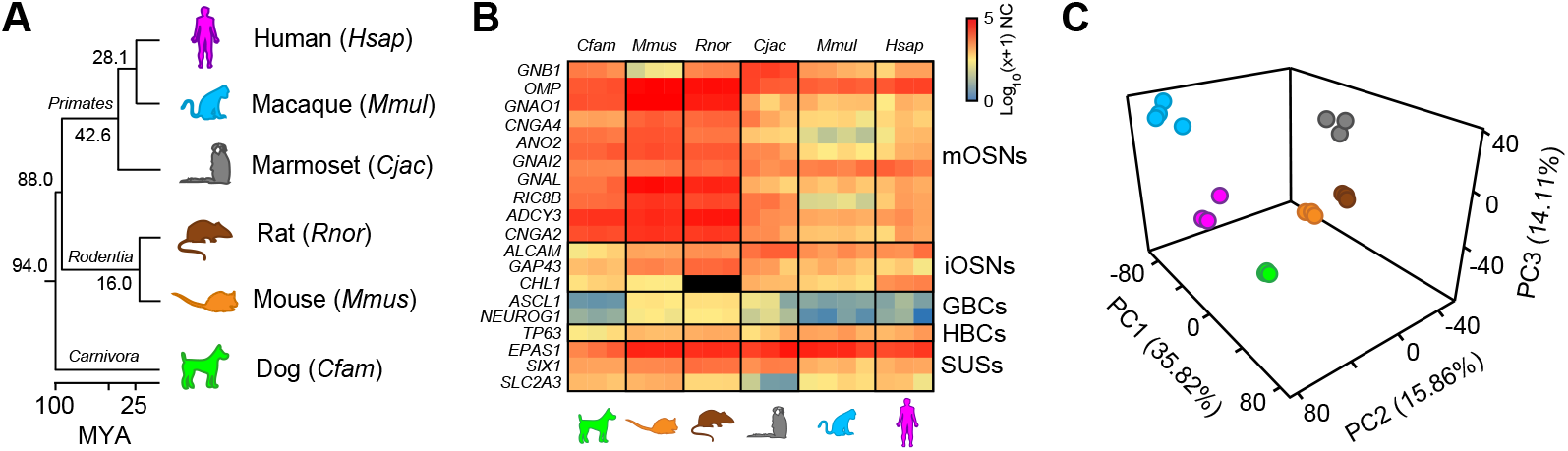
Conservation of WOM expression signatures across mammals. (A) RNA-seq was performed on six species, representing three mammalian lineages: *Carnivora*, *Rodentia*, and *Primates*. Species color scheme used is kept consistent across all figures and is as follows: light green – dog, orange – mouse, brown – rat, dark grey – marmoset, light blue – macaque, fuchsia– human. (B) Heatmap of the expression levels of the canonical markers of the main cell populations present in the WOM samples. RNA expression levels are represented on a log_10_ scale of normalized counts (NC) plus one (0 - not expressed, 5 - highly expressed). There is conservation of expression among all species. mOSNs: mature OSNs, iOSNs: immature OSNs, GBCs: globose basal cells, HBCs: horizontal basal cells, SUSs: sustentacular cells, RES: respiratory epithelium. No *CHL1* ortholog was annotated in the rat genome version analyzed (black squares). (C) Principal component analysis (PCA) of the expression levels for the 9725 ‘one-to-one’ orthologs. Percentages of the variance explained by the principal components (PCs) are indicated in parentheses. PC1 separates rodents from primates, PC2 old-world from new-world primates and hominins, and PC3 separates dog from the other five species.

Gene markers for mature OSNs (mOSNs), immature OSNs (iOSNs), horizontal basal cells (HBCs), and sustentacular cells (SUSs) are highly expressed across all species analyzed (Fig. 1B), consistent with previous studies (Saraiva et al., 2015a; Saraiva et al., 2015b). Genes indicative of globose basal cells (GBCs) are enriched only in rodents and marmoset. Taken together, we have sampled, processed and sequenced RNA of sufficiently high quality to reproducibly capture the neuronal component of WOM from inbred laboratory species (mice and rats) and non-laboratory species (dogs, marmoset, macaques, and humans).

To investigate broader gene expression dynamics of the WOM across mammalian evolution, we focused on the 9725 genes that: i) have 1:1 orthology across the six species, ii) share ≥40% amino-acid identity with the human ortholog, and iii) are expressed in at least three replicates. A principal component analysis (PCA) of the expression data revealed inter-species differentiation among sub-sets of orthologous genes: PC1 primarily separates rodents from primates, PC2 and PC3 separate new-world monkeys (marmosets) from old-world monkeys (macaques) and hominins (humans), and dogs from all the other species, respectively (Fig. 1C and Fig. S1A, B). Hierarchical clustering (HC) analysis further supports these results (Fig. S1C).

Several chemosensory receptors and/or unique molecular barcodes define non-canonical (i.e. not expressing OR genes) OSN subsystems in the WOM (Greer et al., 2016; Hussain et al., 2009; Leinders-Zufall et al., 2007; Liberles and Buck, 2006; Omura and Mombaerts, 2015; Saraiva et al., 2015a; Saraiva et al., 2015b). While orthologs for these marker genes exist in most mammals, it remains unclear whether expression is also conserved in their olfactory systems. We retrieved the annotated orthologs for these genes and analyzed their RNA-seq expression estimates. We found that the WOM of all species expresses trace-amine associated receptors (*TAARs*) and membrane spanning 4-pass A (*MS4As*) chemosensory receptors (Fig. 2A, B). Within each family, the most abundant receptor and relative receptor abundance level varies greatly between species. Although *TAAR5* is the most abundantly expressed gene in human and dog, *TAAR2* is the most expressed in macaque and marmoset, and *Taar6* and *Taar7b* in mouse and rat, respectively (Fig. 2A). Among the *Ms4a* gene family, *MS4A8B* is the most abundant in human, *MS4A14* in macaque, *MS4A7*/*Ms4a7* in marmoset, dog and rat, and *Ms4a6b* in mouse (Fig. 2B). Moreover, our analysis revealed that the expression pattern of the molecular barcode of OSNs expressing guanylate cyclase genes (*GUCY2D/GC-D* or *GUCY1B2*) is also largely conserved (Fig. 2C). In other words, several specialized, non-canonical olfactory sub-systems are likely present across mammals, including human.

**Figure 2.**
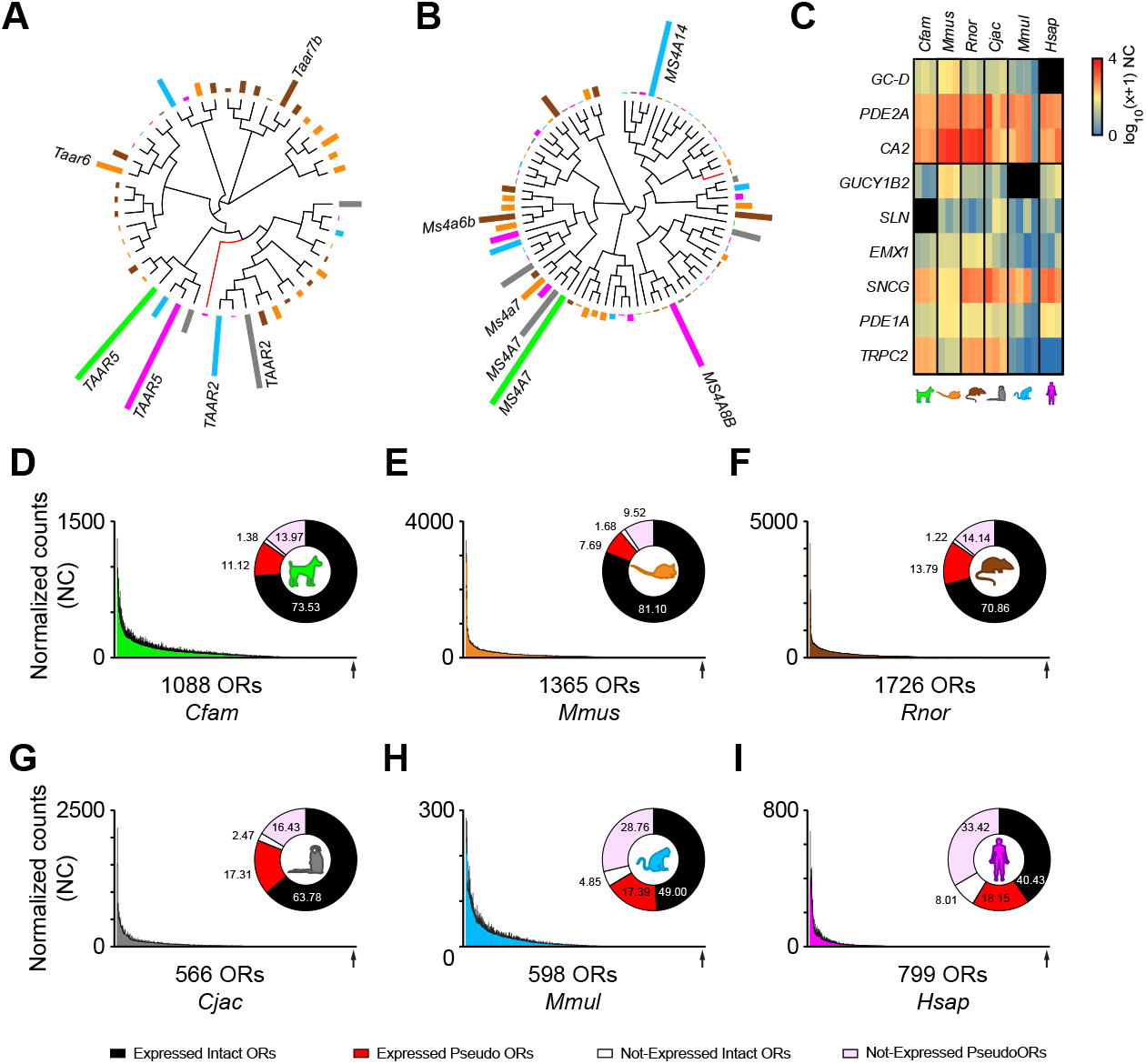
Gene expression profiles of nasal chemosensory receptors in mammals. (A-B) Unrooted phylogenetic tree and mean expression levels for all *TAAR* (A) and *MS4A* (B) receptor orthologs in the six species. Bars indicate the mean contribution (%) of each receptor to the total gene expression within each family, and per species. Red branches indicate pseudo and truncated OR genes. Black branches indicate intact OR genes. (C) Heatmap of the expression pattern for markers of *GC-D*^*+*^ and *Gucy1b2*^*+*^ OSNs in mammals. RNA expression levels are represented on a log_10_ scale of normalized counts (NC) plus one (0-not expressed, 4-highly expressed). *TRPC2* is a pseudogene in human and macaque, and *GUCY1B2* is a pseudogene in human. (D-I) Distribution of mean expression values for each of the OR genes in the WOM of dog (D), mouse (E), rat (F), marmoset (G), macaque (H) and human (I). Genes are displayed in descending order of their mean expression values (normalized counts). Error bars represent the standard error of the mean (SEM) from 3-4 individuals. To the right of each panel, a circular plot shows the percentages of intact and pseudo plus truncated ORs expressed (≥ 1 normalized count) or not expressed (< 1 normalized count) in at least 1 individual. Under the *x*-axis, the total number of ORs within each species, and the position of the least abundant OR (arrow) are noted.

Next, we focused on OR genes, the expression of which are accurate quantitative markers for the abundance of over a thousand OSN subtypes in mouse (Buck and Axel, 1991; Ibarra-Soria et al., 2017). We used uniquely mapped reads to quantify and analyze the distribution of OR-expressing OSN subtypes in the WOM of the six species. Because ORs can be pseudogenes in some individuals but not in others (Griff and Reed, 1995; Mainland et al., 2014), in these analyses we included all ORs annotated as intact and pseudogenes (hereafter referred to as ORs). In these species, the fraction of intact and Class II ORs are higher than the fractions of pseudogenes and Class I OR genes, respectively (Fig S2A). The only exception is human, with a higher fraction (51.56%) of pseudogenes. The OR pseudogene fraction is consistently higher than its relative contribution to the total OR gene expression pool (Fig. S2A), meaning that pseudogenes are, on average, expressed at lower levels than intact genes. A similar pattern emerged when analyzing Class I and Class II ORs (Fig. S2B). Furthermore, we found that expression estimates for ORs are more consistent between replicates from inbred strains (like mouse and rat; r_s_=0.97-0.98, P<0.0001), than replicates from outbred animals (like macaque and human; r_s_=0.68-0.77, P<0.0001, Fig. S2C). Within each species, the percentage of all intact ORs expressed (≥1.0 normalized counts) in at least one replicate varies between 83.46% in human and 98.31% in rat (Fig. 2D-I, File S3). In contrast, between 43.9-58.3% of ORs annotated as truncated or pseudogenes lack expression (<1.0 normalized counts) in any of the replicates (Fig. 2D-I, File S3). Importantly, with this approach we were able to quantify a total of 323 intact human ORs (plus additional 145 truncated or pseudogenes), thus increasing the number of detected intact human ORs detected by ∼18, when compared to a previous study using an array-based approach (Verbeurgt et al., 2014). Together, these results are indicative of the adequacy of our sampling strategy and of the quality of our WOM samples and suggests that most intact ORs are expressed, and putatively functional, in most species. We found that OR genes are expressed across a large dynamic range in all six species, with only a few being highly expressed (Fig. 2D-I). These results are consistent with previous studies in mouse and zebrafish (Ibarra-Soria et al., 2014; Ibarra-Soria et al., 2017; Saraiva et al., 2015a; Saraiva et al., 2015b). As OR expression correlates positively with the number of OSN subtypes in the mouse and zebrafish WOM (Ibarra-Soria et al., 2017; Saraiva et al., 2015a), our results indicate that a few highly abundant OSN subtypes are present in each mammalian species, with the majority present in relatively low numbers.

While the overall distribution pattern of the canonical, or OR-expressing, OSN subtypes is similar among species, the relationship among them remains undetermined. Next, we plotted phylogenetic trees overlaid with the relative frequency of OSN subtypes, as defined by the abundance of the OR gene they express. To compare mean OR expression levels between individuals and species, we transformed the plotted values into the percentage of the total OR expression for each individual (File S3). Species-specific phylogenetic trees revealed that there is no apparent clustering of the ORs that define the most abundant canonical OSN subtypes within each species (Fig. S3A). However, within the order *Rodentia*, mouse and rat display a high level of conservation in OSN subtype frequency (Fig. 3A). In contrast, when comparing abundance levels of OR-expressing OSN subtypes within the order *Primates* (marmoset, macaque, and human) or between any other species combinations, we observe no apparent conservation in OSN subtype representation (Fig. 3B,C). Nonetheless, because of the high sequence similarity and large gene repertoires, it can be difficult to assign 1:1 orthology among all OR genes. To circumvent this problem, we used a classification method that sorts ORs into ortholog gene groups (OGGs) (Niimura et al., 2014). We identified 73 OGGs showing complete 1:1 orthology (Fig. 3D). In other words, each OGG comprises of a set of a single intact orthologous OR in all six species. Of these 73 OGGs, 18 are populated by Class I ORs and 55 by Class II ORs, hereafter referred to as OGG1-and OGG2-, respectively. Hierarchical clustering analysis of OSN subtype abundance divided the 73 OGGs into two major clusters, both containing Class I and Class II ORs. Of the 12 OGGs composing Cluster 1, half (5.67 ±0.80) contain species-specific highly expressed OSN subtypes (i.e., above the 90^th^ percentile) (Fig. 3D), which is significantly above chance level for all species (binomial test, two-tail, P=0.000-0.021, File S4), but marmoset (binomial test, two-tail, P=0.085, File S4). In contrast, the 61 OGGs composing Cluster 2 are populated by OSN subtypes that vary greatly in abundance between species, and of which only ∼9% are highly expressed (Fig. 3D). This is as expected by chance in all species (binomial test, two-tail, P=0.077-0.182, File S4), except human (binomial test, two-tail, P=0.040, File S4). Collectively, these data show that only in a few cases is phylogenetic conservation positively associated with high OR expression, suggesting that it cannot be used as a reliable predictor of highly abundant OSN subtypes in the WOM.

**Figure 3.**
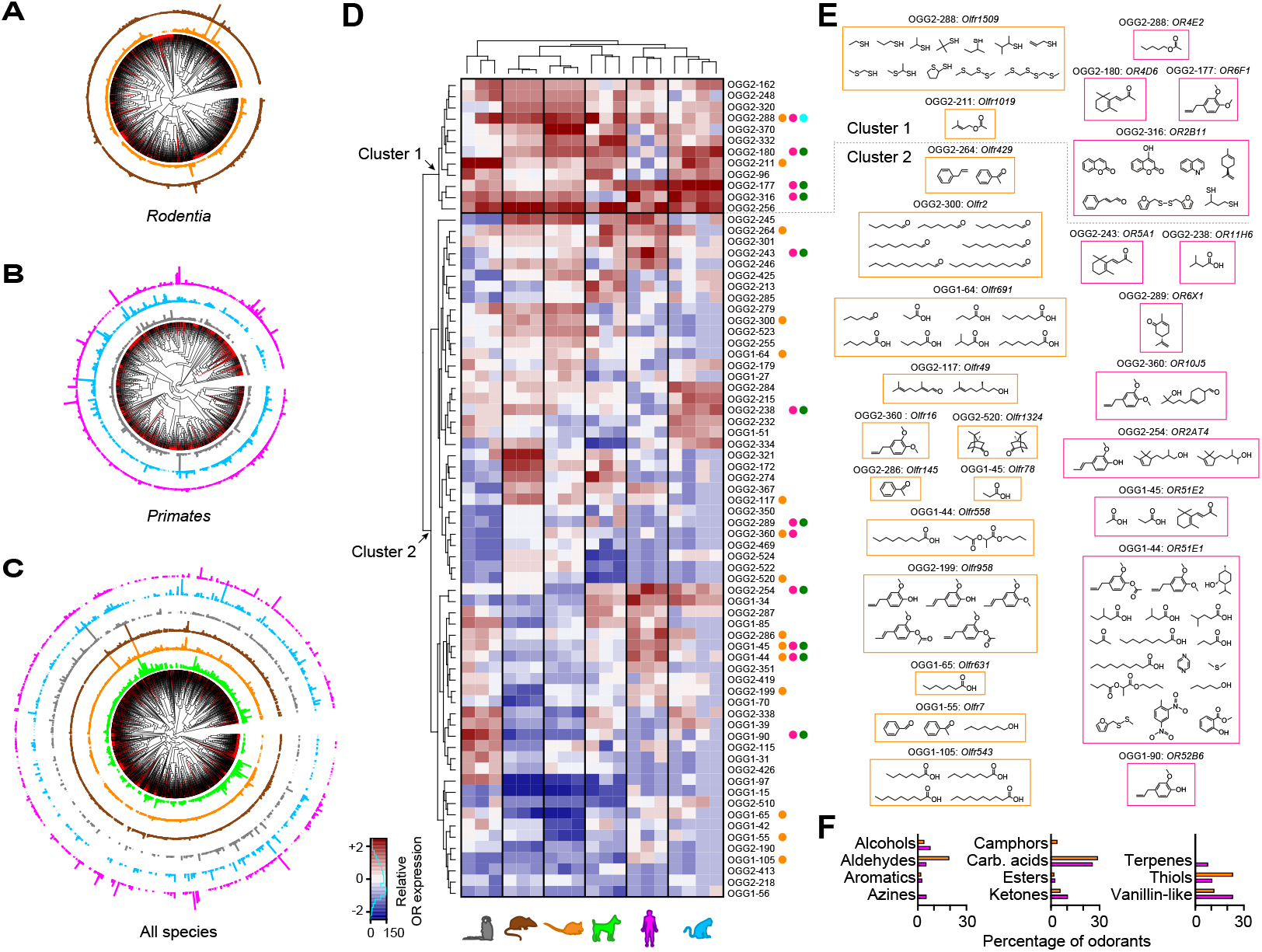
Abundance and ligand biases for highly conserved OR-expressing OSN subtypes across mammalian evolution. (A-C) Unrooted phylogenetic trees containing the mean expression levels for all OR genes from the orders *Rodentia* (A) and *Primates* (B) separately, and for all six species (C). Bars indicate the mean contribution (%) of each receptor to the total gene expression within each receptor family, and per species. Red branches indicate pseudo and truncated OR genes. Black branches indicate intact OR genes. (D) Heatmap of the expression pattern for the ORs populating the highly conserved 73 OGGs across all species. Two clusters were identified (arrows): Cluster 1 containing highly abundant ORs across all species, and Cluster 2 containing highly abundant ORs only in one or a subset of species. Normalized percentage values of expression are represented on a relative abundance scale (-2-lowly abundant, +2-highly abundant). OGGs containing human and/or mouse deorphaned ORs are indicated by orange and fuchsia circles, respectively. Mouse ORs activated by semiochemicals (SMCs) and human ORs activated by key food odorants (KFOs) are indicated by cyan or dark-green circles, respectively. (E) Odorants recognized by the deorphaned mouse (left column and within orange rectangles) and human (right column and within fuchsia rectangles) ORs. In some, but not all cases, ligands for the same OR have similar structures. In 1/3 OGGs (OGG2-288) containing both deorphaned mouse and human ORs, the ligands are different in structure and perceived odors. (F) Percentage of odorants, according to their chemical class, activating mouse and/or human ORs.

RNA-seq expression estimates of chemosensory receptor genes in the WOM are positively correlated with the number of OSNs expressing that given receptor (Ibarra-Soria et al., 2017; Saraiva et al., 2015a). Moreover, increasing the numbers of an OSN subtype may result in increased sensitivity to the odorants that activate these cells (D’Hulst et al., 2016). The interesting hypothesis is raised that for some OSN subtypes, the greater their abundance, the greater their contribution to the perception of its ligand. Under this assumption, unusually highly abundant OSN subtypes could play a role in detecting subsets of odorants that serve a critical ecological function, such as olfactory threshold or hedonics, semiochemical detection, or food choice.

To investigate this hypothesis, we started by identifying ORs above the 90^th^-percentile of expression in each species, hereafter also referred to as “the most” abundant OSN subtype (File S5). By focusing our analysis in ORs with known ligands, we find that ∼36% (26/73) of deorphaned human ORs are above the 90^th^ percentile of expression, which is more than expected by chance (binomial test, two-tail, P<0.0001). In contrast, only ∼15% (13/89) of mouse ORs with known ligands are above the 90^th^ percentile, as expected by chance (binomial test, two-tail, P=0.1552).

We then investigated whether the OSN subtypes expressing ORs in the 73 highly conserved OGGs are more likely to play a role in detecting subsets of related odorants and/or biologically relevant odors. Because most OR-ligand combinations known come from studies in mouse or human, we limited all our downstream analysis to these two species. We found a total of 73 unique odorants that activate mouse and/or human ORs in 23 OGGs, of which 12 contain deorphaned mouse ORs, 8 contain deorphaned human ORs, and 3 contain both deorphaned human and mouse ORs (Fig. 3D,E). These odorants have diverse perceived odors in humans and based on their chemical structures we assigned them to 11 different chemical classes. In both human and mouse, carboxylic acids (cheesy/sweaty odor) are the most represented class (accounting for ∼25% of all detected odorants) (Fig. 3F). Thiols (sulfurous) and vanillin-like (sweet) odorants account for additional 23% in mouse and human, respectively. Interestingly, these two categories together represent ∼44% of all detected odorants in both species. Other categories visibly different between species are aldehydes (fruity, aldehydic) in mouse, and alcohols (floral, fruity) and ketones (fruity, floral, buttery) in human. Interestingly, terpenes (green, minty) and azines (animalic, pungent) were detected only by human, and camphors exclusively by mouse alone. Furthermore, we found that 10/11 OGGs (e.g., OR5A1, OR11H6 and OR6X1) containing deorphaned human ORs that detect molecules (e.g., β-ionone, isovaleric acid and (-)-carvone) defined as human key food odorants (KFOs, Fig. 3D,E and Fig. S3B), which are relevant ecological odorants (Dunkel et al., 2014). In one additional OGG, a mouse OR (*Olfr1509*) is activated by the odorant (methylthio)methanethiol (Fig. 3D,E and S3B), which is known to be a mouse semiochemical (SMC)(Lin et al., 2005). Taken together, these results suggest that ORs displaying high levels of 1:1 orthology across mammals play a critical olfactory role, as they are tuned to recognize specific classes of odorants, which include ecologically relevant odors.

The experiments above revealed a striking feature of the highly conserved, and most abundant mammalian OSN subtypes, and raise two important additional questions: Are highly abundant OSN subtypes within a species biased towards specific odorants? Do the most abundant OSN subtypes in different mammalian species share similar biases? To answer these questions, we first investigated possible biases in the sensory profiles (defined by their human odor qualities) and the physicochemical properties of odorants recognized by the most abundant human and mouse OSN subtypes. Unexpectedly, we found that the sensory profiles of odorants activating exclusively the mouse or human OSN subtypes above the 90^th^ percentile are highly overlapping and highly correlated (r_s_=0.6019, P=0.0227, Fig. 4A). We observed only minor differences between the two species, with humans preferring floral, spicy, dairy and sweaty/pungent smells, and mouse showing a bias towards odorants perceived by humans as nutty, minty/camphor/menthol and sulfurous/meaty odors. PCA analysis of the physicochemical properties of these odorants showed similar results, with all odorants intermingling in the odor space (Fig. 4B).

**Figure 4.**
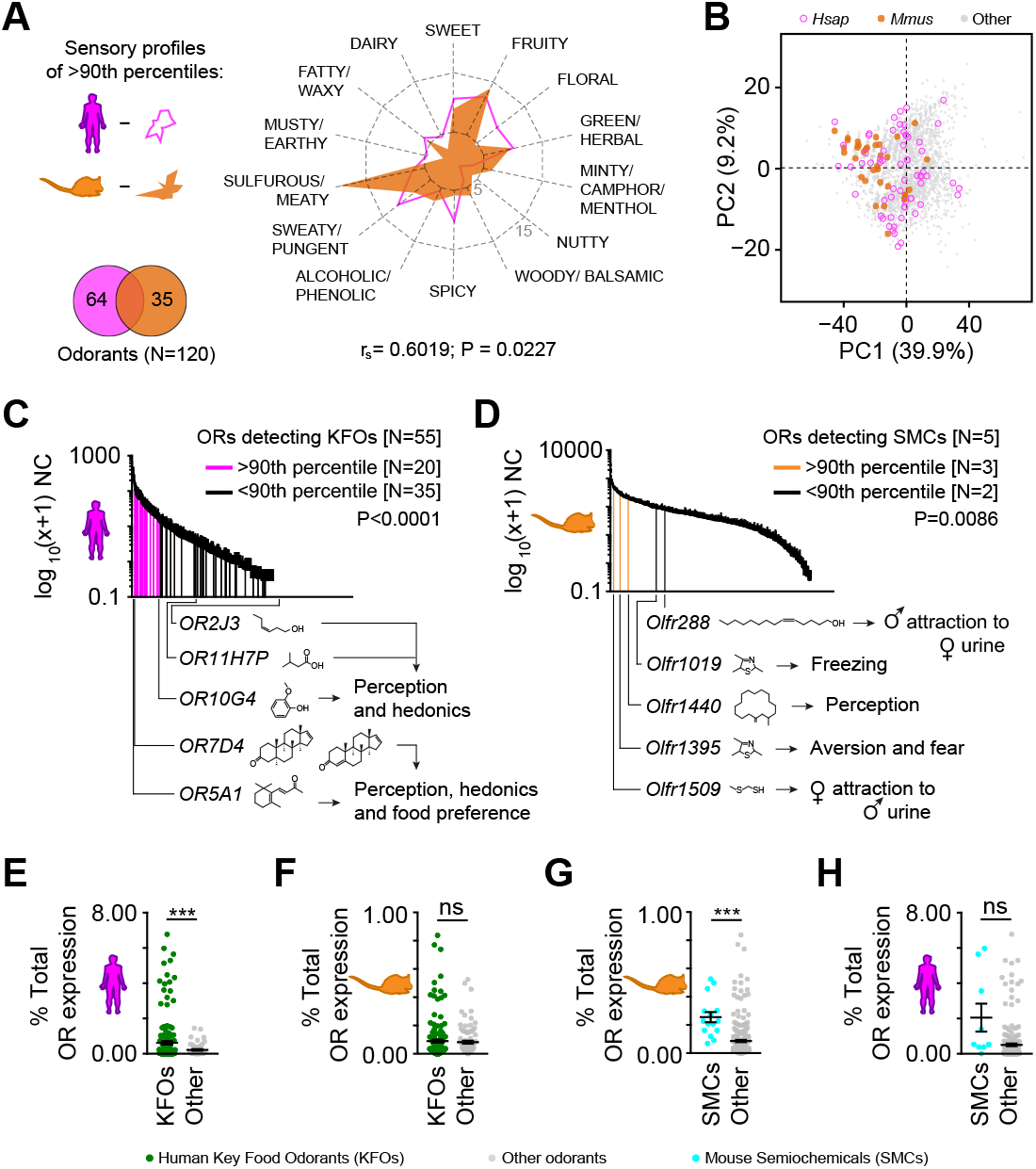
Sensory profile and detection of ecologically relevant odorants by highly abundant ORs/OSN subtypes in mouse and humans. (A) The sensory profiles (spider plot) of the odorants detected exclusively by either mouse or human ORs above the 90^th^-percentile of expression (Venn diagram) are vastly overlapping and significantly positively correlated (r_s_=0.6019, P=0.0227). (B) PCA of the 696 physicochemical descriptors of odorants detected exclusively by either mouse or human ORs above the 90^th^-percentile of expression (see lower left in panel A). Percentages of the variance explained by the PCs are indicated in parentheses. The mouse- and human-specific odorants greatly overlap in the odor space. (C-D) Distribution of mean normalized counts (NC) represented on a log_10_(x+1) scale expression values for each of the OR genes in the human (C) and mouse (D) WOM. OR genes detecting at least one human key food odorant (KFO) or a mouse semiochemical (SMC) are indicated according to their expression percentile (fuschia/orange, above the 90^th^ percentile; black, below the 90^th^ percentile). Known OR-ligand pairs playing a role in olfactory perception, hedonics, or behavior are indicated below the *x-*axis. Error bars represent the standard error of the mean (SEM) from 3 sample replicates. Binomial test, using Wilson/Brown method to calculate the confidence interval (CI), two-tail. (E-F) The mean expression levels of ORs detecting human KFOs are 2.4 times higher in humans (E), but not in mouse (F). Unpaired *t*-test with Welch’s correction, two-tail, ***P≤0.001; ^ns^non-significant (P≥0.05). (G-H) The mean expression levels of ORs detecting mouse SMCs are 2.7 times higher in mouse (G), but not in humans (H). Unpaired *t*-test with Welch’s correction, two-tail, ***P≤0.01, ^ns^non-significant (P≥0.05).

Next, we analyzed the expression levels of all human and mouse OSN subtypes expressing ORs that detect KFOs, SMCs or other odorants. The analysis revealed an enrichment above the 90^th^-percentile of expression for OSN subtypes in both human (binomial test, two-tail, P<0.0001) and mouse (binomial test, two-tail, P= 0.0086) detecting KFOs and SMCs, respectively (Fig. 4C,D). Interestingly, we found that the five human ORs known to contribute to odor perception and hedonics detect KFOs, and the two (*OR7D4* and *OR5A1*) playing a role in food preferences (Jaeger et al., 2013; Keller et al., 2007; Lunde et al., 2012; Mainland et al., 2014; McRae et al., 2012; Menashe et al., 2007) are among the most abundant OSN subtypes (Fig. 4C). Similarly, three mouse ORs (*Olfr1509, Olfr1395* and *Olfr1440*) detecting at least one SMC (Jiang et al., 2015; Lin et al., 2005; Saito et al., 2017; Sato-Akuhara et al., 2016; Yoshikawa et al., 2013) are also above the 90^th^-percentile (Fig. 4D). Intriguingly, OSN subtypes detecting other odorants are also enriched in human (binomial test, two-tail, P= 0.0086) but not in mouse (binomial test, two-tail, P= 0.3730) (Fig. S4A,B), raising the hypothesis that these enrichments could be derived from the fact that highly abundant ORs are more likely to be deorphaned (see results above).

In order to obtain a second line of evidence supporting the putative enrichment (above the 90^th^-percentile of expression) of OSN subtypes sensing human KFOs and mouse SMCs, we compared the mean expression values of these subtypes against the ones binding only other odorants. We found that human ORs detecting at least one KFO have on average ∼2.4-fold higher expression levels than ORs detecting other odorants (unpaired t-test with Welch’s correction, two-tail, P=0.0030; Fig. 4E). Moreover, a similar analysis with mouse ORs revealed no significant differences in expression between OSN subtypes detecting human KFOs or other odorants (Fig. 4F). In line with these results, the average expression levels of mouse ORs detecting at least one SMC is ∼2.7-fold higher than ORs not detecting SMCs (unpaired t-test with Welch’s correction, two-tail, P=0.0004; Fig. 4G). Furthermore, a similar analysis for human revealed no significant differences in expression between ORs detecting mouse SMCs or other odorants (Fig. 4H).

## DISCUSSION

Genes involved in sensing an animal’s immediate environment are under strong evolutionary pressure (Behrens et al., 2014; Hayden et al., 2010; Niimura, 2009; Niimura and Nei, 2005, 2006; Risso et al., 2017; Saraiva and Korsching, 2007; Shichida and Matsuyama, 2009; Young et al., 2010). From identifying kin, food sources and sexually receptive mates to avoiding predation and disease, appropriate perception of environmental sensory cues is critical for survival and reproduction. The importance of sensing the chemical environment is reflected in the genetic investment in encoding ORs, which comprise the largest gene families in mammals. Since each mature OSN expresses just one allele of one receptor gene (Chess et al., 1994; Malnic et al., 2004; Saraiva et al., 2015b), calculating total mRNA abundance of each OR in the WOM samples permitted us to assess which receptors have been favorably selected for expression in OSNs (Ibarra-Soria et al., 2017). Here, we profiled the transcriptomes of the WOM samples from six mammalian species from three different orders (*Carnivora, Rodentia,* and *Primates*), covering ∼95 million years of mammalian evolution.

We first investigated the molecular markers of different cell types composing the WOM. While markers for most major olfactory cell types were identified in the WOM of each of the mammals, they occur at different abundances, possibly reflecting relative differences in cell type composition. These differences are consistent with the variations in total numbers of OSNs across mammalian species, which can range from ∼6-225 million (Horowitz et al., 2014; Kawagishi et al., 2014; Moran et al., 1982; Muller, 1955; Youngentob et al., 1997). As these molecular signatures are already present in the WOM samples from zebrafish (Saraiva et al., 2015a), including the most recently discovered subtype expressing *MS4A* (data not shown), these results are consistent with an ancient origin of all mammalian OSN subtypes, arising at least 450 million years ago.

Conservation was also apparent at the whole transcriptome level. Expression level divergence of highly conserved singleton orthologs between human and other species does not scale with evolutionary time, suggesting a tight regulation of expression of such genes in the mammalian olfactory system. This finding is not unprecedented, as tissue-specific expression profiles can display a wide-spectrum of evolutionary paths (Brawand et al., 2011; Chan et al., 2009; Merkin et al., 2012).

Canonical OR genes are shaped by multiple gene birth and death events, which result in species-specific repertoires with strikingly different numbers of intact genes, pseudogenes, Class I and Class II ORs (Nei et al., 2008). Most intact OR genes show evidence for expression in at least one replicate and most pseudogenes are not expressed, consistent with previous studies (Ibarra-Soria et al., 2014; Ibarra-Soria et al., 2017; Saraiva et al., 2015a; Saraiva et al., 2015b). We observe that the fraction of total OR gene expression originating from pseudogenes is greater for species that are outbred, such as macaque and human. This observation is consistent with greater inter-individual OR genomic variation in outbred species, resulting in a higher fraction of ORs annotated as pseudogenes in the reference genome for which functional alleles are segregating in the population (Griff and Reed, 1995; Mainland et al., 2014; Menashe et al., 2007).

To date, the impact of evolutionary pressures on comparative global OR gene expression across different mammalian species, and hence the corresponding OSN subtype distribution (Ibarra-Soria et al., 2017; Saraiva et al., 2015a), were unknown. We found that the global distribution profile of OSN subtypes expressing ORs is surprisingly similar in the WOM samples from different mammalian species, with the subtypes consistently distributed across a large dynamic range. It is unclear why, in all vertebrate species we have analyzed to date, a few OSN subtypes are present at high frequency with the majority having a low abundance. This distribution pattern may be an emergent property of the complex, multi-step processes that regulate OR singularity (Monahan and Lomvardas, 2015; Tian et al., 2016). The highly abundant OSN subtypes can be consistent between closely related species, like in mouse and rat, but not over greater evolutionary distances. Indeed, we noted above that OR orthology, or the phylogenetic proximity of ORs, is generally a poor predictor of the abundance of the OSN subtypes between species. Together these results suggest the species-specificity of the olfactory sub-genome extends to its regulatory elements. Indeed, ORs expressed at different levels in inbred strains of mice have greater numbers of SNPs upstream of the transcriptional start site compared to those ORs expressed at the same levels. Moreover, these variants impact transcription factor binding sites(Ibarra-Soria et al., 2017).

How does cataloguing the distribution of OSNs in each species advance our understanding of the sense of smell? In humans, loss-of-function variants in the *OR5A1* gene have been shown to increase the odor threshold by about 3 orders of magnitude to its ligand ß-ionone, suggesting that this OSN subtype is the most sensitive to this ligand (Jaeger et al., 2013; McRae et al., 2013). Similarly, mutations in *OR7D4* significantly contribute to specific anosmias of androstenone and androstadienone (Keller et al., 2007). Consistent with this, we observed that OSNs expressing *OR5A1* or *OR7D4* are particularly abundant in human WOM (on average ranked 6^th^ and 11^th^, respectively). Consequently, we propose the more abundant an OSN subtype is, the greater its contribution to the establishment of the detection threshold and perception of its ligands. Genetic variation in other highly expressed ORs may therefore make them strong candidates for causing specific anosmias or hyposmias.

We hypothesize that OSN subtypes that are more abundant in each species may be the consequence of selection for the sensitive detection of ecologically meaningful odorants for that species. Supporting this notion, we found a significant enrichment in abundance for human OSNs that detect KFOs, not observed in mouse OSNs. In contrast we found a similar enrichment for mouse SMC in mouse OSNs, but not human OSNs, albeit with a much smaller sample size.

We propose that the over-representation of specific OSN subtypes is an evolutionary phenomenon: driven by direct selective pressure on genetic elements that promote monogenic OR selection because OSN abundances are, in mouse at least, largely genetically encoded (Ibarra-Soria et al., 2017). However, it remains possible that the distributions of OSN subtypes result from biases in cell survival during neurogenesis, migration, projection or circuit integration. OSN life-span can be altered by odor exposure (Santoro and Dulac, 2012), but when directly compared to the genetic influence the impact of odor environment on OSN abundance is subtle (Ibarra-Soria et al., 2017). Data supporting a model whereby an increase in an OSN subtype abundance driven by odor exposure was trans generationally inherited in mice has been reported (Dias and Ressler, 2014). If this process scaled additively across subsequent generations, our observations could be environmentally driven. However these data have since been challenged on statistical grounds (Francis, 2014). Unless humans differ dramatically from mice in their OSN dynamics, we consider it improbable that human KFO detection in the most abundant OSN subtypes is a physiological response to KFO exposure alone.

We cannot exclude that ascertainment biases contribute to the association between ethologically relevant odorants and high OSN abundance. For example, it is possible that KFOs and SMCs were differentially enriched in odorant libraries screened against orphan ORs in humans and mice respectively. Moreover, in accordance with a previous study (Verbeurgt et al., 2014), our results suggest that ORs expressed in the most abundant OSN subtypes are more likely to be deorphaned than those expressed in the least abundant OSNs, even though mouse and human ORs have been the subject of systematic deorphanisation efforts (Mainland et al., 2015; Saito et al., 2009). Future high-throughput studies focused on unraveling the complete ligand activation patterns of all mammalian ORs will be critical to confirming, or refuting, the conclusions from this study.

## ACKNOWLEDGMENTS

We thank Hiroaki Matsunami and Yoshihito Niimura for insightful discussions.

## AUTHOR CONTRIBUTIONS

L.R.S. initiated the project, designed experiments, performed experiments, interpreted results, supervised the project and wrote the paper; F.R.M.A. performed data analyses; M.M. performed data analyses; E.H.A-M. performed data analyses; C.J.A. performed data analyses; C.T. performed experiments; X.I.S. performed data analyses; M.K. carried out experiments; L.V.G. performed experiments; M. J. performed experiments; M. G. performed experiments; C. O’F. performed experiments; S. M. performed experiments; P.V.J. performed data analyses; P.M. designed experiments; J.C.M. initiated the project, designed experiments; J.D.M. designed experiments and analyzed data; D.W.L. initiated the project, designed experiments, carried out experiments, interpreted results, supervised the project and wrote the paper.

## ADDITIONAL INFORMATION

MG, CO’F and SM were paid employees of Mars Petcare during the period the work was conducted, based at the WALTHAM Centre for Pet Nutrition, and contributed to the analysis of dog ORs only.

