## Supplementary Information for "A transcriptomic atlas of mammalian olfactory mucosae reveals an evolutionary influence on food odor detection in humans"

**MATERIAL AND METHODS**

### Mice

### RNA-sequencing data for the mouse (*Mus musculus*, *Mmul*) WOM samples was retrieved from a previously published study (Ibarra-Soria et al., 2014).

### Rats

### Brown Norway rats (*Rattus norvegicus, Rnor*) were maintained in accordance with UK Home Office regulations, under a project license approved by the Wellcome Trust Sanger Institute Animal Welfare and Ethical Review Body. The entire WOM was collected and immediately frozen and stored at -80^o^C.

### Dogs

Dog (*Canis familiaris, Cfam*) WOM samples were collected from animals submitted by veterinary practices to Hannover University Institute of Pathology for pathological diagnosis. Tissue was collected as soon as possible following euthanasia. The entire WOM was collected before being cut into small sections and snap frozen on liquid nitrogen. Samples were shipped on dry ice and stored at -80^o^C. In all cases owner consent for use of samples in research was obtained.

### Primates

### Rhesus macaques (*Macaca mulatta, Mmul*) and common marmosets (*Callithrix jacchus, Cjac*) were kept at the German Primate Center (Goettingen, Germany). Rhesus macaque WOM samples originate from a study, which was authorized by the governmental veterinary authority (the Lower Saxony State Office for Consumer Protection and Food Safety (Niedersachsisches Landesamt for Verbraucherschutz und Lebensmittelsicherheit LAVES, Ref. No. 33.9-42502-04-14/1456)) according to the regulations of the German Welfare Act (Tierschutzgesetz der Bundesrepublik Deutschland) and the European Directive 2010/63/EU on the protection of animals used for experimental and other scientific purposes. Common marmoset WOM samples originate from animals that were humanely euthanized because of non-infection related animal welfare reasons (e.g. trauma). The use of these samples for this study has been approved by the Animal Welfare and Ethics Committee of the German Primate Center.

### Humans

### Human (*Homo sapiens,* *Hsap*) nasal mucosa/olfactory epithelium was harvested during endoscopic sinus surgery (ESS) for oncological purposes in the University Hospitals of Leuven, Belgium between April 2014 and December 2016. All included patients (N=3) had written informed consent according to the study protocol, approved by the Medical Ethical Committee on Clinical Investigations at the University Hospitals of Leuven on April 23th 2014 (S5648). Included subjects underwent ESS for resection of an adenocarcinoma (stage III-IV) and during the same procedure, olfactory epithelium of the contralateral (healthy) side was harvested at the olfactory groove, followed by post-operative irradiation. After collection of the sample, the tissue was kept in RNA-later and send to the Max Planck Research Unit of Neurogenetics in Frankfurt, Germany, for further analysis.

### RNA-sequencing of Whole Olfactory Mucosa

### RNA from the WOMs was extracted using the RNeasy mini/midi kit (Qiagen), according to the manufacturer’s protocol. mRNA was prepared for sequencing using the TruSeq RNA sample preparation kit (Illumina) with a selected fragment size of 200–500 bp. All samples were sequenced on an Illumina HiSeq 2500, to generate paired-end 100 bp sequencing reads. Libraries generated yielded an average of 49.36 ± 2.61 million (mean ± standard error) reads. Newly generated RNA-seq data from these studies is available in the European Nucleotide Archive (ENA). Accession numbers can be found in File S1.

### RNA-seq Data Processing and Alignment

### To analyze the data, we first created customized GTF annotation files containing all annotated ORs for each mammalian species analyzed in this study. To create custom annotation files for the different species, we downloaded the relevant whole-genome sequences corresponding to previously described OR sequences (Matsui et al., 2010; Niimura et al., 2014; Niimura and Nei, 2007). Genome sequences and Ensembl 79 annotations from mouse (Mus *musculus*; GRCm38), macaque (*Macaca mulatta*; MMul_1) and marmoset (*Callithrix jacchus*; C_jacchus3.2.1) were downloaded from Ensembl (<http://ensembl.org>). Genome sequences and Ensembl 54 annotations from rat (*Rattus norvegicus*; Rnor_4.0), Dog (*Canis familiaris*; CanFam2) and human (*Homo sapiens*; hg18) were downloaded from the UCSC Genome Bioinformatics Site (<http://genome.ucsc.edu>). OR nucleotide sequences were collected via personal communication with relevant authors.

### We generated custom annotated GTF files for use in the RNA-Seq mapping pipeline by first using BLAT (Kent, 2002) to map the OR sequences to their respective genome. Using a custom R script, we removed ORs that mapped with <95% identity to the genome and removed multi-mapping ORs. When 2 ORs overlapped in the same region, the best hit was kept; if there was no best hit, we randomly kept one OR. Using the GenomicRanges (Lawrence et al., 2013) R package we identified overlapping annotations in the downloaded GTF files and replaced these with the OR sequences. Entries for OR sequences that didn’t overlap with any existing annotation were appended thus creating a new GTF file per species.

STAR v 2.4.0i (Dobin et al., 2013) was used to index and map reads to their respective genome using the custom annotations: outFilterType=BySJout, alignSJoverhangMin=8, alignSJDBoverhangMin=1, outFilterMultimapNmax=20, alignIntronMin=20, alignIntronMax=1000000, alignMatesGapMax=1000000, outFilterMismatchNoverLmax=0.04, outFilterMismatchNmax=999. We then performed read summarization using *featureCounts* (Liao et al., 2014). Intra-species normalization of read counts was performed using the DEseq2 (Love et al., 2014) R package. On average, 83.85 ± 1.66 % of the total reads mapped uniquely to the genome.

### OGG cluster assignment and regression analysis

To assess the total number of clusters among the OR genes in the 73 conserved OGGs, we performed hierarchical clustering by creating a dissimilarly matrix based on the normalized percentage value of the expression, assuming the total number of clusters ranging from 2 to 8 (File S4), using the *hclust* function implemented in R. We then compared the stability of the resulting clusters based on cluster statistics including average Silhouette distance (Rousseeuw, 1987), average Pearson gamma (Halkidi et al., 2001), and within-between cluster ratio (a higher value of the former two statistics and a smaller within-between cluster ratio indicates a better fit). Two clusters were determined by balancing the performance of the cluster statistics and total number of clusters for all species. Three cluster for only humans and three cluster for only mice. The R package ‘fpc’ (Hennig, 2015) was used to obtain the cluster statistics.

### Sequence Alignment and Phylogenetic Reconstruction of OR Repertoires

OR protein sequences were aligned using Clustal Omega with default parameters (Li et al., 2015). The sequence alignment was manually edited using Mega 4 (Tamura et al., 2007) as previously described in (Alioto and Ngai, 2005). In brief, sequences were trimmed upstream of the 'G' motif in EC1 (~10 amino acids upstream of the 'GN' motif in TM1) and downstream of the 'K' motif in IC4 (~11 aa downstream of the 'NP' motif in TM7). Positions containing alignment gaps and missing data in most sequences were eliminated. Phylogenetic trees were generated using ClustalW with 100 bootstraps (Li et al., 2015). Visualization and overlay of OR gene expression data was done using Evolview (Zhang et al., 2012).

**Odorant information and odor descriptors**

### All odorant structures and associated CAS numbers were retrieved from either Sigma-Aldrich (<http://www.sigmaaldrich.com/>) or Pubchem ([https://pubchem.ncbi.nlm.nih.gov/](https://pubchem.ncbi.nlm.nih.gov/))). Odor descriptors were retrieved using the GoodScents Company database (<http://www.thegoodscentscompany.com/)>.

### A comprehensive list of the cognate mouse and human OR-ligand pairs was assembled (last update: June 2017) by combining the data contained in the ODORactor database with literature searches (Duan et al., 2012; Dunkel et al., 2014; Geithe et al., 2016; Liu et al., 2011; Mainland et al., 2014; Mainland et al., 2015; Noe et al., 2016; Saito et al., 2009; Verbeurgt et al., 2014). Human key food odorants (KFOs) and mouse semiochemicals (SMCs) were identified from databases and previously published studies (Duan et al., 2012; Dunkel et al., 2014; Jiang et al., 2015; Keller et al., 2007; Kobayakawa et al., 2007; Lin et al., 2005; Lunde et al., 2012; Mainland et al., 2014; McRae et al., 2012; Menashe et al., 2007; Saito et al., 2017; Sato-Akuhara et al., 2016; Takahashi et al., 2005; Verbeurgt et al., 2014; Yoshikawa et al., 2013). Additionally, we found that the human OR detecting muscone (*OR5AN1*), also recognizes the structurally related key food odorant ß-ionone (Figure S4).

**PCA analysis with the physico-chemical descriptors**

### We calculated 4,885 physicochemical descriptors for 2,662 molecules using Dragon 6.0 (Talete) software. We removed descriptors where >90% of the values were identical, where the most common value was >19x more common than the second most common value, or where values were missing for any odor. The remaining 696 descriptors were then used in principal component analysis on the 2,662 compounds to plot the 113 molecules of interest in the context of the full set.

### Luciferase assay

### In vitro activity of the human *OR5AN1* was measured using the Dual-Glo Luceriferase Assay System (Promega). Hana3A cells were co-transfected with the olfactory receptor, a short from of receptor transporter protein 1 (RTP1S), the type 2 muscarinic acetylcholine receptor (M3-R), Renilla luciferase driven by an SV40 promoter, and firefly luciferase driven by a cyclic AMP response element. 18-24 hours post-transfection, ORs were treated with medium or serial dilutions of odorants spanning 1nM to 1mM in triplicate. Odors were first diluted to 1M stocks in DMSO, then diluted from stocks to the appropriate concentration in CD293 (Gibco). Four hours after odorant stimulation, luciferase activity was measured using the Synergy 2 (BioTek). Normalized luciferase activity was calculated by dividing firefly luciferase values by Renilla luciferase values for each well. Results represent mean response (for 3 wells) +/- s.e.m. Responses were fit to a three-parameter sigmoidal curve.

### Statistical Analysis

Statistical analyses were done using GraphPad Prism (version 6.04), PAlaeontological STatistics (version 3.14, <http://folk.uio.no/ohammer/past/>), and the R statistical language. Data values were standardized and hierarchical clustering analysis was performed using Euclidean distances with Ward’s method. For principal component analysis, the data matrix was standardized and correlation matrixes used to compute the eigenvalues and eigenvectors.

**SUPPLEMENTARY FIGURE LEGENDS**

**Figure S1 - Conservation of the WOM expression signatures across mammals**

(A, B) Principal component analysis (PCA) of the expression levels for the 9725 orthologs. Percentages of the variance explained by the principal components (PCs) are indicated in parentheses. PC1 separates rodents from primates, PC2 old- from new-world primates, and PC3 separates dogs from the remaining species.

(C) Hierarchical clustering analysis (HC) of the tissue expression profiles for the 9725 Biomart ortholog pairs. Bootstrap values (100 boostraps, 1 represents > 0.999) for the 5 major nodes are indicated.

**Figure S2 – OR gene expression in mammals**

(A, B) Percentages of Intact/Pseudogenes (A) and Class I/ Class II (B) OR genes in the genomes of the 6 analyzed mammalian species. NC – normalized counts. Note: the ‘Pseudogenes’ category includes both truncated and pseudogenes.

(C) Intraspecific spearman correlation coefficients for OR repertoires are higher between replicates from inbred strains (i.e. mouse and rat), than replicates from outbred animals (i.e. dog, marmoset, macaque and human).

**Figure S3 – Abundance and ligand biases for highly conserved canonical/OR-expressing OSN subtypes across mammalian evolution**

(A) Unrooted phylogenetic trees containing the mean expression levels for all ORs for all six species analyzed. Bars indicate the mean contribution (%) of each receptor to the total gene expression within each receptor family, and per species. Red branches indicate pseudo and truncated OR genes. Black branches indicate intact OR genes.

(B) Hierarchical clustering analysis of the expression pattern for the ORs populating the highly conserved 73 OGGs across all species. OGGs containing human and/or mouse deorphaned ORs are indicated by orange and fuchsia circles, respectively. Mouse or human ORs activated by semiochemicals (SMCs) or key food odorants (KFOs) are indicated by cyan or dark-green circles, respectively.

**Figure S4 – Distribution of mouse and human OR genes that detect exclusively other odorants.**

(A, B) Distribution of mean normalized counts (NC) represented on a log10(x+1) scale expression values for each of the OR genes in the human (A) and mouse (B) WOM. OR genes detecting only other odorants are indicated according to their expression percentile (fuschia/orange, above the 90th percentile; black, below the 90th percentile). Error bars represent the standard error of the mean (SEM) from 3 sample replicates. Binomial test, using Wilson/Brown method to calculate the confidence interval (CI), two-tail.

**Figure S5 – The muscone human olfactory receptor, *OR5AN1*, is also activated by the key food odorant β-ionone.**

Hana3A cells were co-transfected with expression vectors encoding *OR5AN1* (or vector alone), a short from of receptor transporter protein 1 (RTP1S), the type 2 muscarinic acetylcholine receptor (M3-R), Renilla luciferase driven by an SV40 promoter, and firefly luciferase driven by a cyclic AMP response element. ORs were treated with medium or serial dilutions of odorants spanning 1nM to 1mM in triplicate. Odors were first diluted to 1M stocks in DMSO, then diluted from stocks to the appropriate concentration in CD293 (Gibco). Normalized luciferase activity was calculated by dividing firefly luciferase values by Renilla luciferase values for each well. Results represent mean response (for 3 wells) +/- s.e.m. Responses were fit to a three-parameter sigmoidal curve.

**File S1 – Sample information, accession numbers and RNA-seq quality metrics.**

A dataset containing detailed information about each sample processed for RNA-seq, its ENA accession numbers and several quality control (QC) metrics.

**File S2 – WOM expression estimates for dog, mouse, rat, marmoset, macaque and human.**

A dataset containing the expression values (normalized counts) for all genes in the WOM of the six analyzed mammals.

**File S3 – Expression estimates for the OR repertoires of dog, mouse, rat, marmoset, macaque and human.**

A dataset containing the summary statistics of the ORs expressed (≥ 1 normalized count) or not expressed (< 1 normalized count) in at least 1 individual, and the expression values (normalized counts) for all ORs in the WOM of the six analyzed mammals.

**File S4 – Composition of the highly conserved 73 OGGs across mammals**

A dataset containing the summary statistics, OGG composition, cluster number solution and the hypergeometric test results regarding the highly conserved 73 OGGs across all species.

**File S5 – Expression estimates of the most and least abundant ORs or OSN subtypes.**

A dataset containing the all ORs above the 90th- and below the 10th-percentile of expression for each analyzed species. These have also been referred throughout the text as “the most” and “the least” abundant ORs or OSN subtypes, respectively (File S6).
