## Supplementary figures and images for "A transcriptomic atlas of mammalian olfactory mucosae reveals an evolutionary influence on food odor detection in humans"

### Figure S1

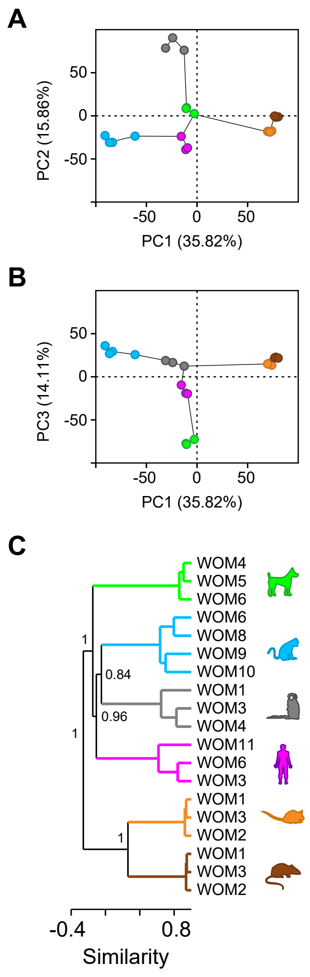

### Figure S2

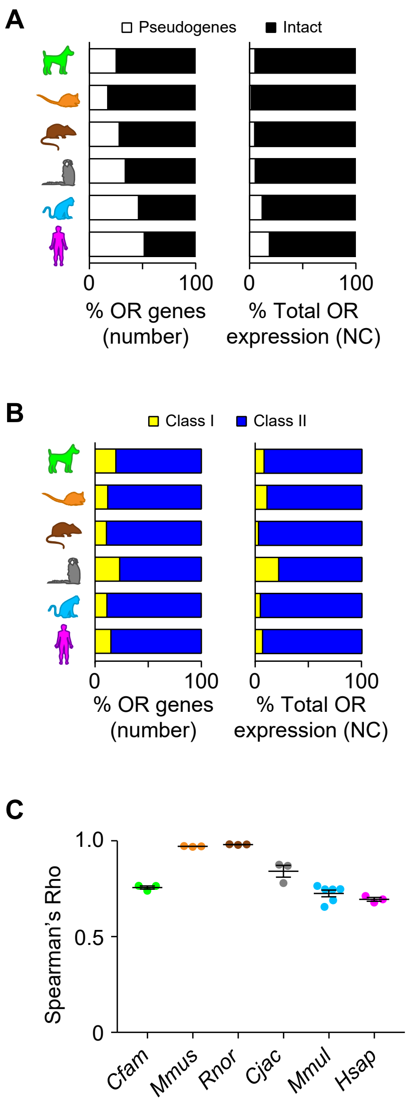

### Figure S3

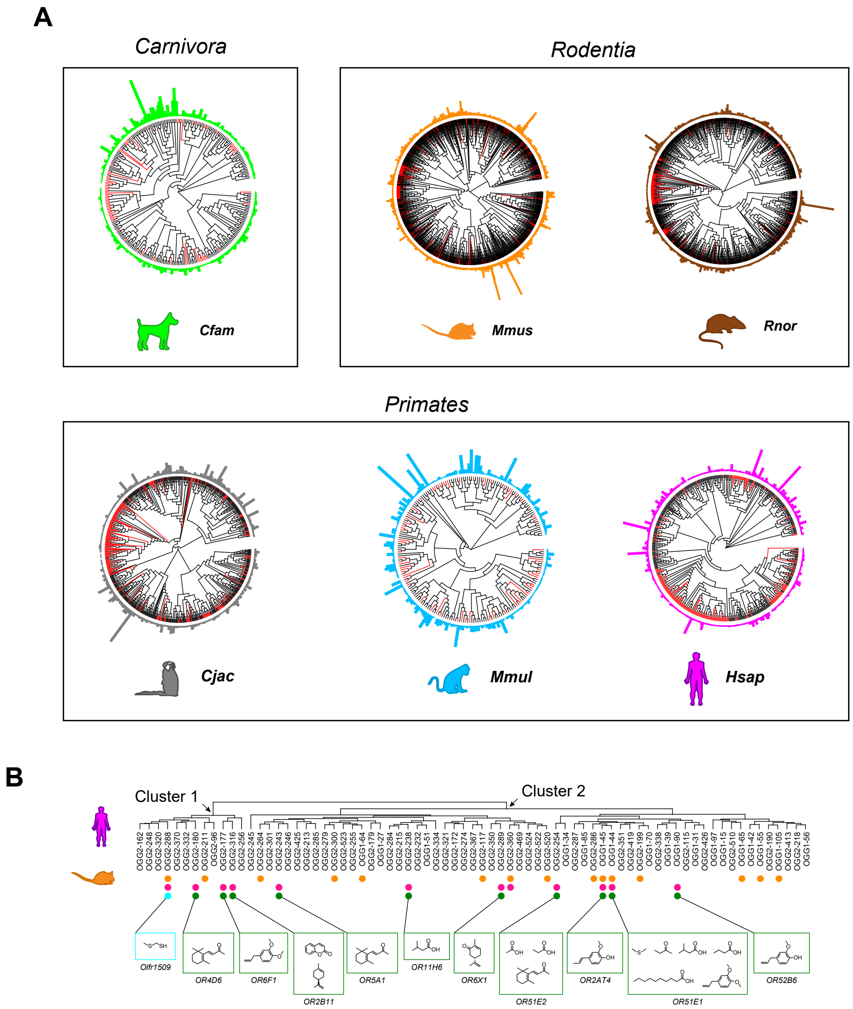

### Figure S4

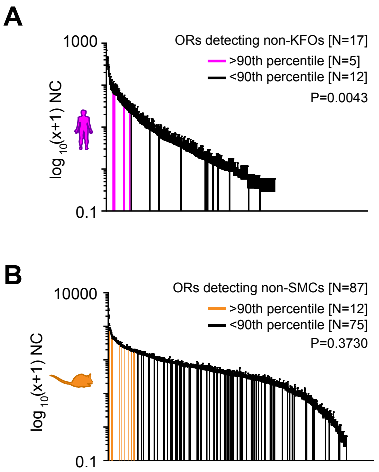

### Figure S5

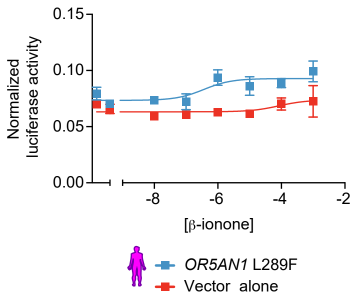
